# Discovery of three novel neutralizing antibody epitopes on the human astrovirus capsid spike and mechanistic insights into virus neutralization

**DOI:** 10.1101/2024.09.14.613010

**Authors:** Sarah Lanning, Nayeli Aguilar-Hernández, Vitor Hugo B Serrão, Tomás López, Sara M. O’Rourke, Adam Lentz, Lena Ricemeyer, Rafaela Espinosa, Susana López, Carlos F. Arias, Rebecca M. DuBois

## Abstract

Human astroviruses (HAstVs) are a leading cause of viral childhood diarrhea that infect nearly every individual during their lifetime. Although human astroviruses are highly prevalent, no approved vaccine currently exists. Antibody responses appear to play an important role in protection from HAstV infection, however knowledge about the neutralizing epitope landscape is lacking, as only 3 neutralizing antibody epitopes have previously been determined. Here, we structurally define the epitopes of 3 uncharacterized HAstV-neutralizing monoclonal antibodies: antibody 4B6 with X-ray crystallography to 2.67 Å, and antibodies 3H4 and 3B4 simultaneously with single-particle cryogenic-electron microscopy to 3.33 Å. We assess the epitope locations relative to conserved regions on the capsid spike and find that while antibodies 4B6 and 3B4 target the upper variable loop regions of the HAstV spike protein, antibody 3H4 targets a novel region near the base of the spike that is more conserved. Additionally, we found that all 3 antibodies bind with high affinity, and they compete with receptor FcRn binding to the capsid spike. These studies inform which regions of the HAstV capsid can be targeted by monoclonal antibody therapies and could aid in rational vaccine design.

**Importance:** Human astroviruses infect nearly every child in the world, causing diarrhea, vomiting, and fever. Despite the prevalence of human astroviruses, little is known about how antibodies block virus infection. Here, we determined high-resolution structures of the astrovirus capsid protein in complex with three virus-neutralizing antibodies. The antibodies bind distinct sites on the capsid spike domain. We find that the antibodies block virus attachment to human cells and prevent capsid spike interaction with the human neonatal Fc receptor. These findings support the use of the human astrovirus capsid spike as an antigen in a vaccine to prevent astrovirus disease.

## Introduction

Human astroviruses (HAstVs) are a significant cause of childhood viral diarrhea worldwide, with over 35% of children having had a HAstV infection by age 2.^1^ These small nonenveloped RNA viruses are typically spread by fecal-oral or salivary routes.^2,3^. While HAstV infections are typically self-limiting in immunocompetent people, they can persist as a chronic infection in immunocompromised individuals.^4,5,6^ Young children and immunocompromised individuals are the populations most at risk for HAstV disease, particularly in lower-income or tropical countries where higher burdens of diarrheal disease and additional comorbidities may exist.^7,8^ The classical HAstV clade includes eight serotypes (HAstV1-8), with serotype 1 being the most prevalent worldwide ^9,10,11^ Divergent VA and MLB clades, which may have arisen from animal astroviruses, have been found to cause fatal encephalitis in immunocompromised individuals, and additionally, there has been a report of central nervous system involvement by classical HAstV.^12,13^ Encephalitic symptoms can also be caused by some animal astroviruses, such as mink and bovine astroviruses.^14,15^ Notably, astrovirus-associated encephalitis has been found to be endemic in mink and pig farms, where animals are maintained under intensive production conditions.^16^ Despite HAstV’s prevalence and global health impacts, there are currently no vaccines or HAstV-specific therapeutics available.

However, the development of vaccines against HAstV seems feasible, since some evidence suggests the presence of lasting HAstV immunity induced by prior infection.^1^ Seroprevalence to HAstV in adults is very high (>90%)^17,18^, and HAstV disease is rarer in adults than in children.^19^ Additional studies have supported that the presence of anti-HAstV antibodies may help to protect from severe HAstV disease,^3^ and one case study showed improvement in a patient with chronic HAstV disease after immunoglobulin therapy. However, the mechanism of how antibodies neutralize HAstV is not well understood, in part due to a lack of knowledge surrounding how antibodies interact with the viral capsid, and which parts of the exposed viral capsid are critical for its function.

The HAstV virion consists of a small proteinaceous icosahedral capsid roughly ∼40 nm in diameter, which shelters a ∼7 kb single-stranded positive sense polyadenylated RNA genome. The immature capsid is made up of 180 units of capsid protein originating from open reading frame 2 (ORF2), and displays a T=3 symmetry. The capsid protein is initially expressed as a 90 kDa protein (VP90), which undergoes an intracellular caspase cleavage that is important for viral release from the cell, resulting in a ∼70 kDa (VP70) protein after the loss of its C-terminal acidic domain.^20,21^ In this state, the virus remains immature and must undergo further extracellular protease cleavage(s) to reach its mature infectious form. The exact extracellular protease used *in vivo* for this cleavage event is unknown, but *in vitro* cleavage with trypsin results in a 10^5^ fold increase in infectivity.^22^ This extracellular protease cleavage event cleaves VP70 into the core domain (VP34), and spike domain (VP25/VP27), and additionally removes 60 of the initial 90 dimeric spikes along 5-fold symmetry axes, resulting in 30 dimeric spikes (VP27) remaining on the mature capsid along the 2-fold symmetry axes.^23,24^ The spike domain is known to be important for attachment and entry of the virus, and antibodies that target the spike domain have been found to neutralize HAstV in cell culture, whereas antibodies that target the core domain have not been reported to neutralize HAstV.^25,26^ Recently, two preprint articles reported the identification of the neonatal Fc receptor (FcRn) as an important host receptor for human astrovirus entry, and FcRn was found to bind the HAstV capsid spike.^27,28^ However, information about which regions of the spike are important for this interaction and how antibodies may interfere with this function remains mostly unexplored.

Only three HAstV-neutralizing antibody epitopes have been structurally defined, revealing two neutralizing antigenic sites on the HAstV spike, since two of the neutralizing antibodies (3E8, PL-2) have overlapping epitopes.^29,30^ Both neutralizing epitope regions were located around a conserved putative receptor binding site on the surface of the HAstV spike, known as the “P-site,” and these antibodies were additionally shown to block spike attachment to cells.^29^ Whether these antibodies prevent FcRn binding or some other host factor interaction remains unknown.

Furthermore, whether additional neutralizing antigenic sites exist on the HAstV spike remains unknown. Here, we structurally define three novel neutralizing antibody epitopes, assess their epitope location relative to conserved regions of the HAstV spike, and provide evidence supporting their mechanism of HAstV neutralization.

## Materials and Methods

### Cells and viruses

Caco-2 cells, clone C2Bbe1 (ATCC), were propagated in high-glucose Dulbecco’s modified Eagle’s medium (DMEM-HG) (Sigma) supplemented with nonessential amino acids (Gibco) and 15% fetal bovine serum (FBS) (Cansera) in a 10% CO_2_ atmosphere at 37°C. HAstV serotypes 1 and 2 have been described previously.^26^ All viral strains were activated with trypsin and grown as described before.^31^

### Expression and purification of recombinant HAstV1 and HAstV2 capsid spike proteins

Recombinant HAstV1 and HAstV2 spikes were produced as described previously.^32,31^ Briefly, cDNA corresponding to HAstV1 capsid protein residues 429 to 645 (GenBank: AAC34717.1) or HAstV2 Oxford strain capsid protein residues 429 to 644 (GenBank: KY964327.1) were cloned into pET52B with a C-terminal thrombin cleavage site and a 10-histidine purification tag sequence. Recombinant spikes were expressed in *Escherichia coli* BL21(DE3) and purified from soluble lysates by HisTrap metal-affinity chromatography. Purified HAstV spikes were dialyzed into TBS (10 mM Tris pH 8.0 and 150 mM NaCl).

### Expression and purification of recombinant monoclonal antibody Fabs 3B4, 3H4, and 4B6

The protein-coding sequence of antibodies 3H4, 3B4, and 4B6 heavy and light chains were determined as described previously.^33^ The protein-coding sequences of 3H4, 3B4, and 4B6 light chain and the 3H4, 3B4, and 4B6 heavy chain antigen-binding fragment (Fab) were cloned into separate pCMV plasmids in-frame with an N-terminal human IgG1 signal sequence. The Fab heavy chains were cloned in-frame with a C-terminal thrombin-cleavable double StrepTagll affinity tag. A total of 120 µg of heavy chain plasmid and light chain plasmid combined were added to 8 x 10^7^ CHO-S cells in an OC-400 cuvette (MaxCyte) and were electroporated. CHO-S cells were resuspended in CD-OptiCHO media (Gibco: #12681029) and fed CHO feed (CHO CD EfficientFeed A (Gibco: #A1023401) supplemented with 7 mM L-glutamine, 5.5% glucose, and 23.4 g/L yeastolate) every 24 hours. CHO-S cells were given a final concentration of 1 mM sodium butyrate and maintained at 32 °C, 8 % CO_2_, 85% humidity, 135 rpm, 24 hours after electroporation for 8-10 days. CHO-S cells were centrifuged, and the resulting supernatants were given 1X protease inhibitor cocktail (Millipore 539137), BioLock (Iba Lifesciences 2-0205-050) to block free biotin in the media, and Strep Wash Buffer (50 mM Tris pH 7.4, 150 mM NaCl, 1mM EDTA), and were 0.22 µm filtered. Samples were loaded onto a regenerated StrepTrap HP 5 ml column (Cytiva), washed with Strep Wash Buffer, and eluted with an increasing linear gradient of Strep Elution Buffer (Strep Wash Buffer + 2.5 mM desthiobiotin).

### Expression and purification of recombinant monoclonal antibody scFv 3B4, 3H4, 4B6

Codon-optimized cDNA encoding the 4B6 variable heavy chain and variable light chain connected by a GGS(GGGGS)_3_ linker, were cloned into a derivative pCDNA3.1 vector in frame with an N-terminal human IgG1 signal sequence and a C-terminal thrombin- cleavable double StrepTagll affinity tag. A total of 120 µg of this plasmid was added to 8 x 10^7^ CHO-S cells and was electroporated. scFv 4B6 expression and purification were performed as described above for Fabs. Purified scFv was dialyzed into TBS. Synthetic genes for scFv 3H4 and 3B4 constructs were designed with the light and heavy chain variable domains connected by a GGS(GGGGS)_3_ linker and flanking BgIII and NheI restriction sites. The gene was codon optimized for *Drosophila melanogaster* and ordered from Integrated DNA Technologies. The gene was cloned into a pMT_puro_BiP vector via restriction digest in frame with an N-terminal BiP secretion signal and a C-terminal thrombin cleavable double StrepII affinity tag in the vector. pMT-puro_BiP vectors containing scFv 3H4 and scFv 3B4 were used to make stably transfected Schneider 2 (S2) cells as described previously.^29^ Expression and purification was performed as described previously. ^29^

### Expression and purification of recombinant neonatal Fc receptor (FcRn)

Codon-optimized cDNA encoding the ectodomain of the FCGRT gene (UniProt: P55899, Met1-Ser297) or the β-2-Microglobulin gene (UniProt: P61769, Met1-Met119) were cloned separately into a derivative pCDNA3.1 vector.^34^ The FCGRT construct also contained a C-terminal thrombin-cleavable double StrepTagll affinity tag. A total of 40 µg of FCGRT plasmid and 80 µg of β-2-Microglobulin plasmid were added to 8 x 10^7^ CHO-S cells and electroporated. CHO cell expression was performed as described above. The supernatant was loaded onto a StrepTrap XT affinity column (Cytiva), washed with Strep Wash Buffer, and eluted with Elution Buffer (Strep Wash containing 50mM biotin). Purified FcRn was dialyzed into TBS.

### Binding assays of HAstV in the presence of neutralizing antibodies

Serial 1:5 dilutions of the ascitic fluids for 3B4, 3H4, or for 4B6, were pre-incubated with infectious, purified HAstV1 or HAstV2 particles, respectively (multiplicity of infection [MOI] = 30), for 1 h at room temperature. Caco-2 cell monolayers grown in 48-well plates were washed once with PBS, and then blocking solution (1% BSA in PBS) was added for 45 min at room temperature, followed by a 15 min incubation on ice. The cells were then washed once with ice-cold PBS and incubated with the virus-antibody complex for 1 h on ice. MAb 2D9, which neutralizes HAstV8, was used as a negative control. The unbound virus was washed three times with cold PBS, and the total RNA was extracted with TRIzol Reagent (Invitrogen) according to the manufacturer’s instructions. Viral RNA or cellular 18S RNA was reverse transcribed using MMLV reverse transcriptase (Invitrogen). RT-qPCR was performed with the premixed reagent Real Q Plus Master Mix Green (Amplicon), and the PCR was carried out in an ABI Prism 7500 Detection System (Applied Biosystems). The primers used to detect HAstV1 RNA were Fwd 5’ -ATGAATTATTTTGATACTGAAGAAAATTACTTGGAA - 3’ and Rev 5’ - CTGAAGTACTTTGGTACCTATTTCTTAAGAAAG - 3’. For detection of HAstV2 RNA were Fwd 5’ -ATGAATTATTTTGATACTGAAGAAAGTTATTTGGAA - 3’ and Rev 5’ - CTGAAGTACTGTGGTACCTATTTCTTAAGAAAG - 3’. For normalization, 18S ribosomal cellular RNA was amplified and quantified using forward primer 5’- CGAAAGCATTTGCCAAGAAT - 3’ and reverse primer 5’ - GCATCGTTTATGGTCGGAAC - 3’.

### Assay to determine if the neutralizing antibodies detach HAstV particles bound to cells

Confluent Caco-2 cell monolayers in 48-well plates were blocked with 1% BSA in PBS for 45 min at room temperature followed by a 15 min incubation on ice. Purified HAstV-1 or HAstV-2 particles were added at an MOI of 30 and then incubated for 1 h on ice to allow the binding of the virus to the cell surface. The unbound virus was subsequently removed by washing three times with cold PBS. Serial 1:5 dilutions of the indicated ascitic fluids of either 3B4 or 3H4 for HAstV1, or 4B6 for HAstV2 were added to the cells and then incubated for 1 h on ice. After this incubation, the antibody and unbound virus were removed with cold PBS, and RNA extraction and RT-qPCR quantification were performed as described above. MAb 2D9, which neutralizes HAstV8, was used as a negative control.

### X-ray crystallography structure determination of HAstV2 spike/scFv 4B6 complex

Thrombin digestion was used to remove the Histidine-tag from HAstV2 spike and to remove the StrepII tag from scFv 4B6 (10 U thrombin/mg of protein incubated at 4 °C on a rotating plane overnight). Digestion of StrepII tag from scFv 4B6 and Histidine-tag from HAstV2 spike was confirmed by SDS-PAGE where no visually detectable undigested product was observed. HAstV2 spike was incubated with 2X molar excess scFv 4B6 per spike monomer and the resulting complex was purified by size-exclusion chromatography on a Superdex 75 10/300 GL column. Fractions corresponding to HAstV2 spike/scFv 4B6 complex were determined by peak comparison with gel filtration standards (peak elution volume corresponding to ∼100 kDa) and SDS-PAGE analysis.

Fractions of purified HAstV2 spike/scFv 4B6 complex were pooled and concentrated to 5 mg/ml in TBS pH 8.5. HAstV2-spike/scFv 4B6 protein crystals were formed in 2 µl drops containing a 1:1 ratio of protein solution to well solution consisting of 0.1 M Tris-HCl pH 8.5, and 0.74 M sodium citrate pH 5.5, using hanging drop vapor diffusion at 22 °C. A single crystal was transferred into a cryoprotectant solution consisting of well solution and 18% glycerol, and was then flash-frozen into liquid nitrogen. The Advanced Photon Source synchrotron beamline 23-ID-D was used to collect a diffraction dataset with wavelength 1.0332 Å at cryogenic temperatures. The dataset was processed and scaled using DIALS (ccp4i2)^35^ with a resolution cutoff of 2.67 Å based upon CC_1/2_ and I/σl statistics. A trimmed model of HAstV2 spike (PDB: 3QSQ) and a trimmed model of scFv 4B6 generated by SWISS-model using tremelimumab Fab as a template (PDB: 5GGU) was used for molecular replacement with Phaser. The structure was then manually modeled using Coot^36^ and refined in Phenix.^37^ The final model was deposited into the Protein Data bank (PDB 9CN2).

### Single-particle CryoEM structure determination of HAstV1 spike/Fab 3B4/Fab 3H4 complex

Thrombin digestion was used to remove the Histidine-tag from HAstV1 spike and remove the StrepII tags from Fab 3H4 and Fab 3B4 as described above. HAstV1 spike was complexed with 2X molar excess Fab 3B4 and the resulting complex was purified by SEC on Superdex 200 10/300 GL column. Fractions corresponding to the HAstV1 spike/Fab 3B4 complex were determined by peak comparison with molecular weight standards (peak elution volume corresponding to ∼100 kDa) and SDS-PAGE analysis. These fractions were pooled, and the resulting complex was then mixed with 1.5X molar excess Fab 3H4 and purified by SEC on a Superdex 200 10/300 GL column. Fractions corresponding to the full HAstV1 spike/Fab 3B4/Fab 3H4 complex were determined by peak comparison with gel filtration standards (peak elution volume corresponding to ∼200 kDa) and SDS-PAGE analysis. Fractions of purified HAstV1 spike/Fab 3H4/Fab 3B4 complex were pooled and concentrated to 0.86 mg/ml in 10 mM Tris pH 7.0 and 150 mM NaCl. 3 µl of protein complex was mixed with 0.5 µl of 25 µM lauryl maltose neopentyl glycol (LMNG) detergent to remove orientation bias and was then deposited onto glow discharged UltrAuFoil R.12/1.3 gold grids 400 mesh, blotted using a ThermoFisher Scientific (TFS) Vitrobot Mark IV at 4 °C and 100% humidity, and then plunge frozen into liquid ethane. Grids were screened at UCSC’s Biomolecular CryoEM facility using a TFS Glacios 200 kV microscope coupled to a Gatan K2 Summit direct detector. The top-selected grids were then sent to the Pacific Northwest Center for Cryo-EM (PNCC #160263) for data collection on a TFS Krios G3i microscope coupled to a Gatan K3 Biocontinuum Gif.

7,235 movies containing 60 frames each were collected using a pixel size of 0.415 Å/pixel in super-resolution mode (105,000 x) and an electron dose of 32.26 e/A^2^. Movies were preprocessed (motion correction and CTF estimation) in CryoSPARC v4.3.2.^38^ Initial particle identification was performed using an unbiased blob picker, resulting in 4,132,753 particles, further extracted in a box size 686 pixels. After multiple rounds of 2D classification, 55 top-selected classes containing 214,273 particles underwent the *Ab-initio* reconstruction.

3 selected volumes were generated and then 3D-classified and further refined. The best-representing 3D class was used to create 2D references for a round of template picking. 2,718,470 particles were extracted with a box size 686 pixels, and then underwent on the similar previously established workflow. The top 72 classes containing 262,500 particles were used in a new *Ab-initio* reconstruction. The 2 generated classes were 3D-classified, where one class resulted in an overall gold-standard resolution (FSC_0.143_) of 5.70 Å and containing 138,147 particles. This volume was selected and underwent non-uniform refinement, and further non-uniform refinement using a mask encompassing the entire particle, resulting in a 3D reconstructed volume at 3.74 Å. Unused particles were added from the previous 2D classification, and all particles received local CTF refinement, resulting in a volume of 3.43 Å and 163,237 particles after non-uniform refinement. Additional rounds of local CTF refinement were performed and a mask in which the constant domains of the Fab were removed was used to align particles in local refinement, in order to improve the tridimensional alignment and local resolution of the epitope regions, resulting in the final reconstructed map at 3.33 Å overall resolution. The sharpened map (B factor -112 Å^2^) was opened in ChimeraX (version 1.5.0) and starting models of the HAstV1 spike (PDB: 5EWO) and AlphaFold 3 models of Fabs 3H4 and 3B4 were fitted into the volume. Since the Fab constant domain volume density was poor, the constant domains were removed from the models. The initial model representing the complex was opened in Coot (version 0.9.1) and underwent several rounds of manual refinement and global real-space refinement and was validated using Phenix and MolProbity. The final reconstructed map was deposited in the Electron Microscopy Data Bank (EMD-45427) and the final model was deposited into the Protein Data Bank (PDB: 9CBN). The raw data was made available in EMPIAR DOI: https://doi.org/10.6019/EMPIAR-12182.

### Biolayer interferometry K_D_ determination of neutralizing antibodies 3B4, 3H4, and 4B6

Biolayer interferometry assays on an Octet RED384 instrument were used to determine binding affinity dissociation constants (K_D_). Assays were performed in Octet Kinetics Buffer (PBS pH 7.4 + 0.1% BSA + 0.02% Tween 20) for Fabs 3H4 and 3B4, or Octet Kinetics Buffer + biocytin (PBS pH 7.4 + 0.1% BSA + 0.02% Tween 20 + 50 µM biocytin) for Fab 4B6. For assays with Fabs 3H4 and 3B4, pre-equilibrated Anti-Penta- His (HIS1K) biosensor tips were dipped into Octet Kinetics Buffer for 60 seconds for an initial baseline reading, dipped into 0.5 µg/ml histidine-tagged HAstV1 spike diluted in Octet Kinetics Buffer for 180 seconds to load the sensor tip, and dipped into Octet Kinetics Buffer for 60 seconds for a second baseline reading. Biosensors were then dipped into 4-point serial dilutions of Fab in Octet Kinetics Buffer, consisting of 2.5 nM, 5 nM, 10 nM, and 20 nM for Fab 3H4, and 20 nM, 40 nM, 80 nM, 160 nM for Fab 3B4.

This association step was run for 180 seconds, and then biosensors were dipped into Octet Kinetics Buffer to measure dissociation for a total of 600 seconds. For assays with Fab 4B6, pre-equilibrated Anti-Penta-His (HIS1K) biosensor tips were dipped into Octet Kinetics Buffer + biocytin for 60 seconds for an initial baseline reading, dipped into 0.5 µg/ml histidine-tagged HAstV2 spike diluted in Octet Kinetics Buffer + biocytin for 180 seconds to load the sensor tip, and dipped into Octet Kinetics Buffer for 60 seconds for a second baseline reading. Biosensors were then dipped into a 4-point serial dilutions of Fab 4B6 in Octet Kinetics Buffer, consisting of 25 nM, 50 nM, 100 nM, and 200 nM. This association step was run for 60 seconds, and then biosensors were dipped into Octet Kinetics Buffer to measure dissociation for a total of 60 seconds. These shorter association and dissociation steps were chosen due to the lower affinity of the Fab 4B6.

Kinetics data was processed the same way for all Fabs in the Data Analysis HT software. The baseline step was used to align traces and apply inter-step correction. A reference sample well containing only a spike-loaded biosensor dipped into no analyte (Fab) was subtracted. Savitzky-Golay filtering was used on the traces. For curve fitting, a 1:1 model was globally applied to the dilution series, and fit was evaluated based on R^2^ and χ^2^ values and visual inspection. Average K_D_ values are reported as the average of the three replicates.

### Biolayer interferometry competition assay of Fabs versus FcRn for HAstV spike

Biolayer interferometry competition assays were performed with an Octet RED384. Pre-equilibrated Streptavidin (SA) biosensors tips were dipped into Octet Kinetics Buffer for 60 seconds for an initial baseline reading. For Fabs 3H4 and 3B4, 0.5 µg/ml of biotinylated HAstV1 spike was loaded onto SA biosensors tips for 300 seconds, and dipped into Octet Kinetics Buffer for 30 seconds for a baseline reading. Biosensors were then dipped into either 150 nM Fab 3H4 or 250 nM Fab 3B4 in Octet Kinetics Buffer for 600 seconds to ensure saturation of all spike binding sites, dipped into Octet Kinetics Buffer for 30 seconds as a baseline reading, and then dipped into 2 µM FcRn in Octet Kinetics Buffer for 300 seconds. For Fab 4B6, the assay was performed with the same methods, but 0.5 µg/ml biotinylated HAstV2 spike was used during the antigen loading step, Octet Kinetics buffer + biocytin was used for all assay steps after antigen loading, 250 nM Fab 4B6 was using during the antibody association step, and due to its lower affinity 250 nM of Fab 4B6 was also included in the baseline step after antibody association and in the FcRn sample to maintain saturation of Fab 4B6 on spike. All assays contained additional controls such as a sample in which the primary Fab was dipped into the same concentration of self Fab instead of FcRn to ensure that saturation was achieved, and also a control in which FcRn was bound to the HAstV spike in the absence of any Fab in the first association. All assays were performed in duplicate.

Competition data was processed the same way for all Fabs in the Data Analysis HT software in the Epitope Binning module. A matrix representing competition was generated using the shift between the last 10% average of the signal from the second association and the last 10% average of the signal from the primary association step. The signal from the control in which primary Fab was dipped into self Fab for the second association was subtracted in the matrix row (the shift values from all samples containing the respective Fab), such that full competition is represented by “0.” The signal from the control sample in which FcRn was associated to HAstV spike with no Fab in the primary association was used to normalize in the matrix column (the shift values from samples with FcRn binding in the secondary association step) such that maximum FcRn binding with no competition represents “1.” This normalization is done separately for Fab 4B6 vs Fab 3H4 and 3B4 assays given the different spike serotypes to which FcRn is associated to, but is displayed in the same table. Fabs were considered to compete with FcRn if FcRn binding in the presence of Fab was reduced by 50% or more (a value of 0.5 or lower). The values shown are an average of duplicate assays.

### scFv 3B4, scFv 3H4, and 4B6 neutralization assays

The indicated concentration of antibody or scFv was preincubated with HAstV1 (for scFv 3H4, scFv 3B4 and mAb 3B4) or HAstV2 (for scFv 4B6) at an MOI of 0.02, for 1 h at room temperature. The virus-antibody mixture was then added to confluent Caco-2 cell monolayers grown in 96-well plates and incubated for 1 h at 37°C. After this time, the cells were washed three times with minimum essential medium (MEM) without serum, and the infection was left to proceed for 18 h at 37°C. Infected cells were detected by an immunoperoxidase focus-forming assay, as described previously.^26^

### Data availability

Coordinates and structure factors for the HAstV2 spike/scFv 4B6 complex structure was deposited in the Protein Data Bank (www.rcsb.org) under accession code 9CN2. For the HAstV1 spike/Fab 3B4/Fab 3H4 complex structure, the final reconstructed map was deposited in the Electron Microscopy Data Bank (EMD-45427) and the final model was deposited into the Protein Data Bank under accession code 9CBN. The raw data was made available in EMPIAR at https://doi.org/10.6019/EMPIAR-12182.

## Results

### HAstV-neutralizing antibodies 3B4, 3H4, and 4B6 bind with high affinity to the HAstV spike

We previously generated a panel of IgG1 monoclonal antibodies (mAbs), three of which were found to neutralize either HAstV1 (mAbs 3B4 and 3H4) or HAstV2 (mAb 4B6) in Caco-2 cells, the gold standard cell line used for HAstV propagation and infectivity studies. Here, we generated recombinant antigen-binding fragments (Fabs) of these HAstV-neutralizing antibodies to remove the avidity effects of a full bivalent mAb given the homodimeric nature of their target, the HAstV capsid spike domain. To determine binding affinities, biosensors loaded with HAstV spike were dipped into serial dilutions of Fabs. All three Fabs bind the corresponding HAstV spike with high affinities, with dissociation constants (K_D_s) in the mid-low nanomolar range (Table 1, Supp. Fig. 1).

**Table 1:** Antibodies 3B4, 3H4 and 4B6 bind HAstV spike with high affinity.

| Antibody to spike | Average $K_D \pm \sigma$ (nM) | $\chi^2$ | $R^2$ |
| --- | --- | --- | --- |
| Fab 3H4 - HAstV1 spike | $0.490 \pm 0.002$ nM | <0.2339 | >0.9991 |
| Fab 3B4 – HAstV1 spike | $11.8 \pm 0.5$ nM | <0.7474 | >0.9909 |
| Fab 4B6 – HAstV2 spike | $161 \text{ nM} \pm 2 \text{ nM}$ | <0.0675 | >0.9943 |

These results indicate that immunization with recombinant HAstV spikes are able to induce high affinity HAstV-neutralizing antibodies in mice. Interestingly, Fab 4B6 has the lowest affinity of the three antibodies, yet has the most potent neutralizing activity.^26^

This observation may indicate that other factors besides affinity, such as the location of the antibody binding site or avidity, may influence HAstV neutralization, or this result could be related to serotype difference.

### HAstV-neutralizing antibodies 3B4, 3H4, and 4B6 block attachment of HAstV to Caco-2 cells

To further investigate the mechanism of antibody neutralization, we tested whether mAbs 3B4, 3H4, or 4B6 could block attachment of HAstV to Caco-2 cells, and whether these antibodies could detach the virus which was already bound to cells. Caco-2 monolayers were incubated with HAstV-antibody complexes, or HAstV alone. Unbound virus was washed away and the bound virus was quantified using RT-qPCR. We found that all three antibodies were able to block virus attachment to cells in a dose-dependent manner compared to a negative control antibody (Fig. 1). Interestingly, only 4B6 was able to detach pre-bound virus (Fig. 2). This detachment does not appear to be a function of a high affinity displacement, as 4B6 had the lowest affinity of the three antibodies (Table 1). This could indicate that other factors, such as the binding location, may play a role in an antibody’s ability to detach virus. It is also possible that the serotype affects the ability of the virus to be displaced, as 3H4 and 3B4 neutralize serotype 1, while 4B6 neutralizes serotype 2. The inability of 3H4 and 3B4 to detach virus is also in contrast to previously characterized antibodies 2D9 and 3E8 which neutralize serotype 8 and were able to detach bound virus.^26^

**Figure 1:**
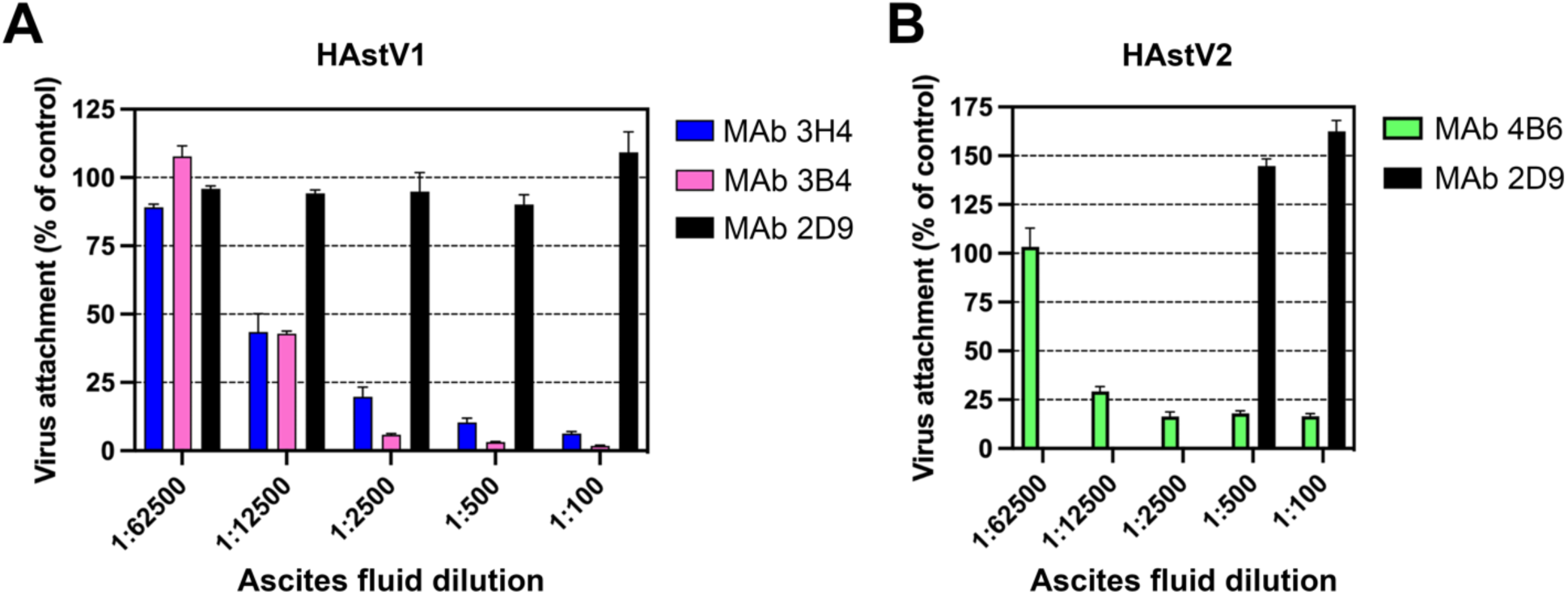
Monoclonal antibodies to HAstV1 and HAstV2 block attachment of the virus to Caco-2 cells. Ascitic fluid of (A) mAbs 3B4 or 3H4 to HAstV1 or (B) mAb 4B6 to HAstV2 block attachment when pre-incubated with the corresponding virus before cell adsorption. MAb 2D9 which is specific to serotype HAstV8 was used as a negative control. Experiments were performed on ice to prevent virus endocytosis. The assay was performed in biological quintuplicates and carried out in duplicate. The data are expressed as percentages of the virus attached in the absence of antibodies and represent the mean ± SEM.

**Figure 2.**
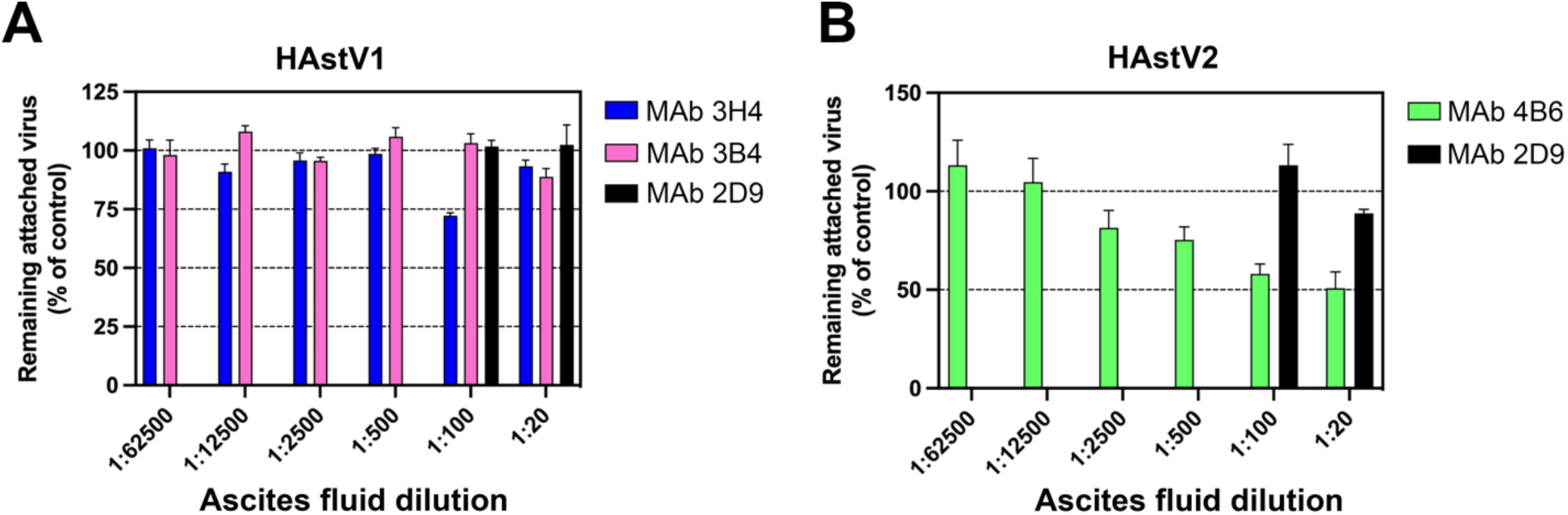
Monoclonal antibody detachment of HAstV previously bound to the surface of Caco-2 cells. (A) HAstV1 or (B) HAstV2 was attached to cells on ice to prevent virus endocytosis. Subsequently, either ascitic fluid of (A) mAb 3B4 or 3H4, or (B) mAb 4B6 was added to cells and incubated for 1 h on ice. The remaining attached virus was determined by RT-qPCR. MAb 2D9 which is specific to serotype HAstV8 was used as a negative control. The assay was performed in biological sextuplicates and carried out in duplicate. The data are expressed as percentages of the virus that remained attached in the absence of antibodies and represent the mean ± SEM.

### HAstV-neutralizing antibodies 3B4, 3H4, and 4B6 compete with FcRn binding to HAstV spike

With the recent discovery of FcRn as a critical receptor for HAstV infection, we investigated whether the HAstV-neutralizing antibodies 3B4, 3H4, and 4B6 could compete with FcRn’s ability to bind to the HAstV spike. Either HAstV1 or HAstV2 spikes were loaded onto biosensors and then dipped into saturating levels of Fab 3H4, 3B4, or 4B6. The biosensors were then dipped into FcRn and these binding shifts were compared to the binding shifts of FcRn to spike-loaded biosensors in the absence of Fab. From this assay, we determined that Fabs 3H4 and 4B6 fully block FcRn binding (Table 2), suggesting that these Fabs either directly or sterically block FcRn’s ability to bind the spike protein. Fab 3B4 does not appear to fully block FcRn binding, but reduces FcRn binding to 40% of the control. Given that 3B4 is still efficient at neutralizing HAstV1, Fab 3B4 may have an alternative mechanism of neutralizing HAstV, such as blocking the interaction of another putative receptor, or the full-length mAb may be necessary for full steric hindrance of the FcRn interaction with HAstV1 spike. These data suggest that one mechanism of antibody neutralization may be by blocking the FcRn interaction with the HAstV spike.

**Table 2:** Antibodies 3B4, 3H4, and 4B6 compete with FcRn receptor to HAstV spike.

|  | FcRn |
| --- | --- |
| Fab 3B4 | 0.394 |
| Fab 3H4 | 0.035 |
| Fab 4B6 | 0.063 |
| no Fab | 1.000 |

**Table 3:**
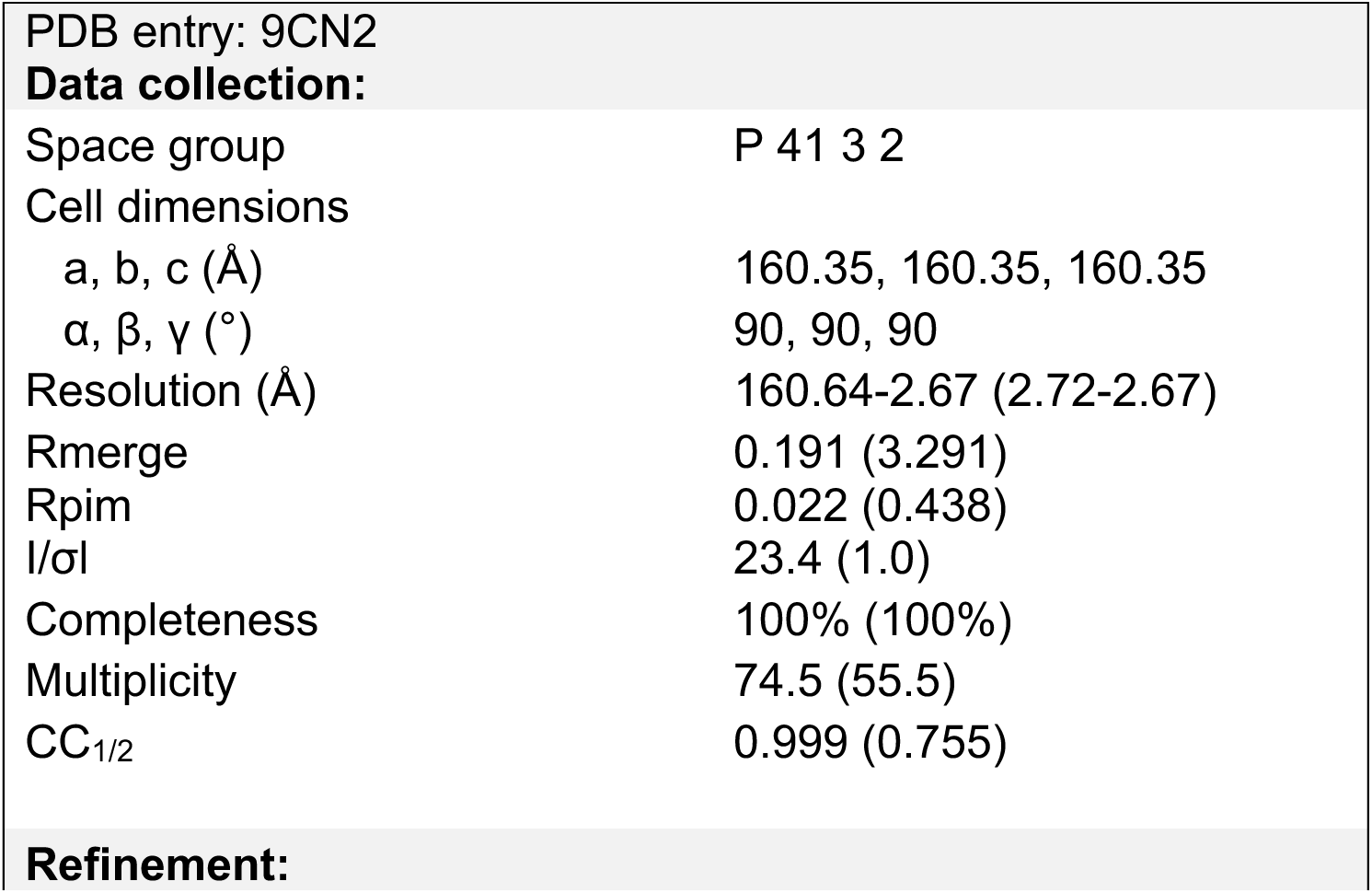
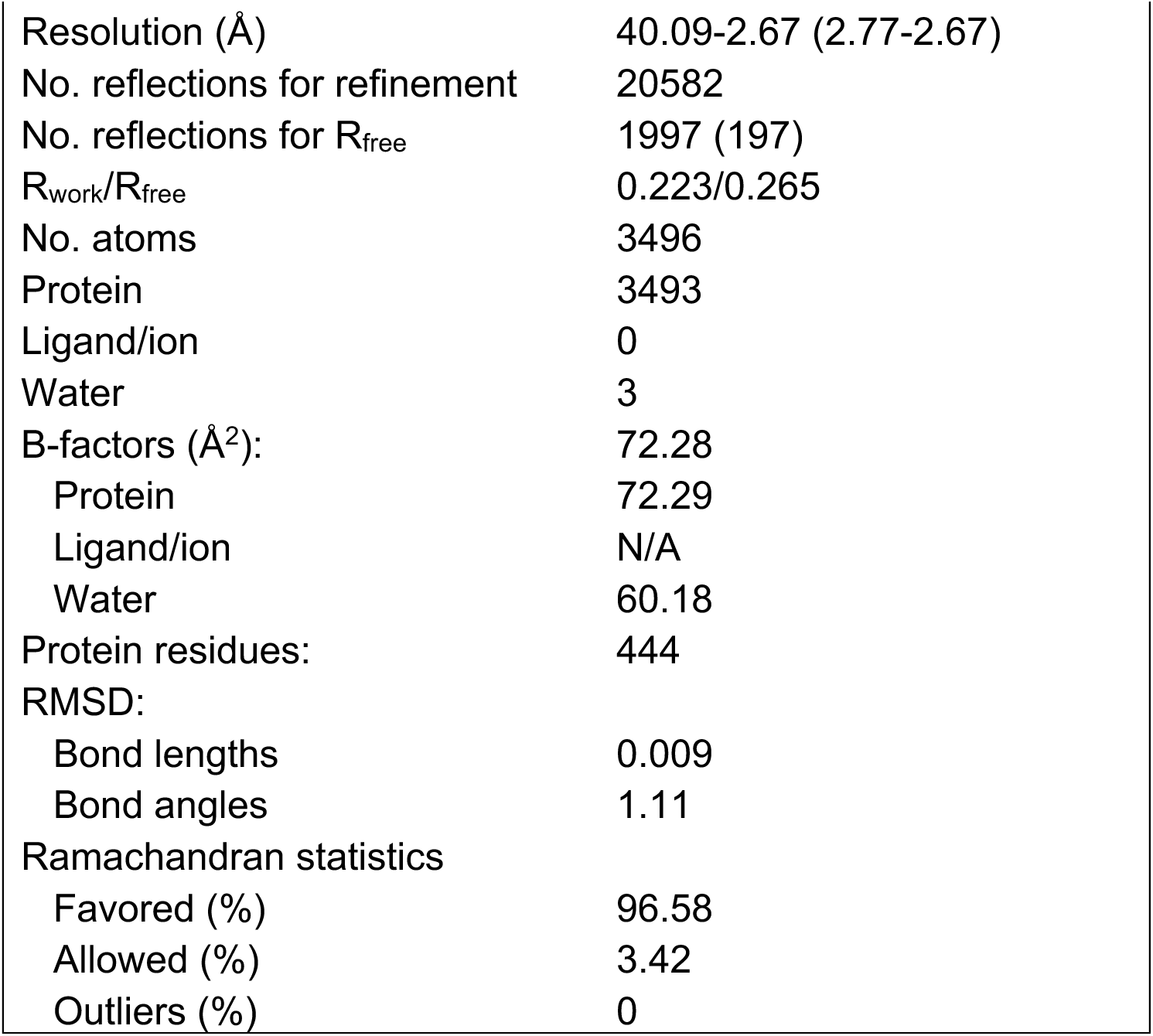
Crystallographic statistics for scFv 4B6 / HAstV2 spike complex.

**Table 4:**
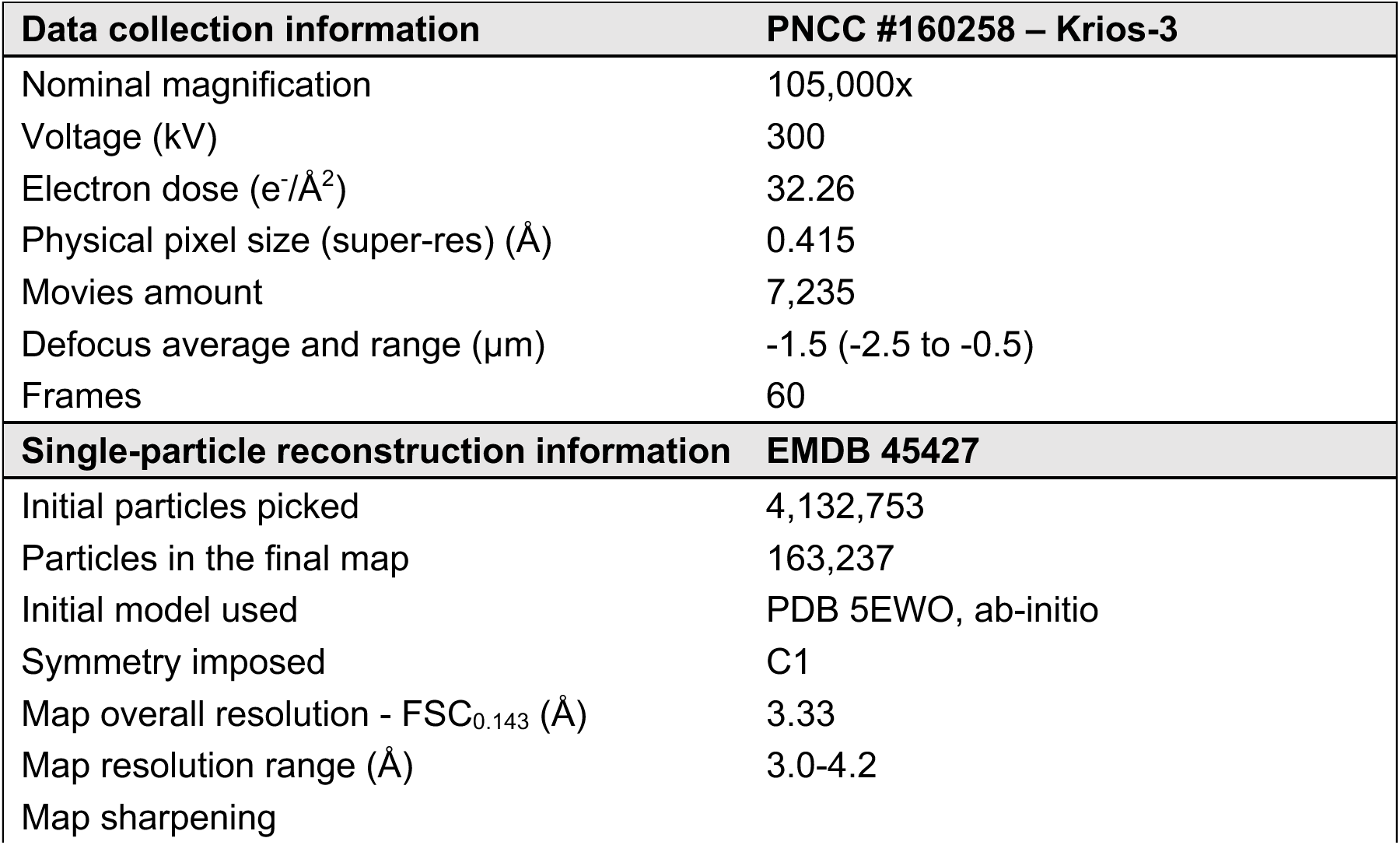
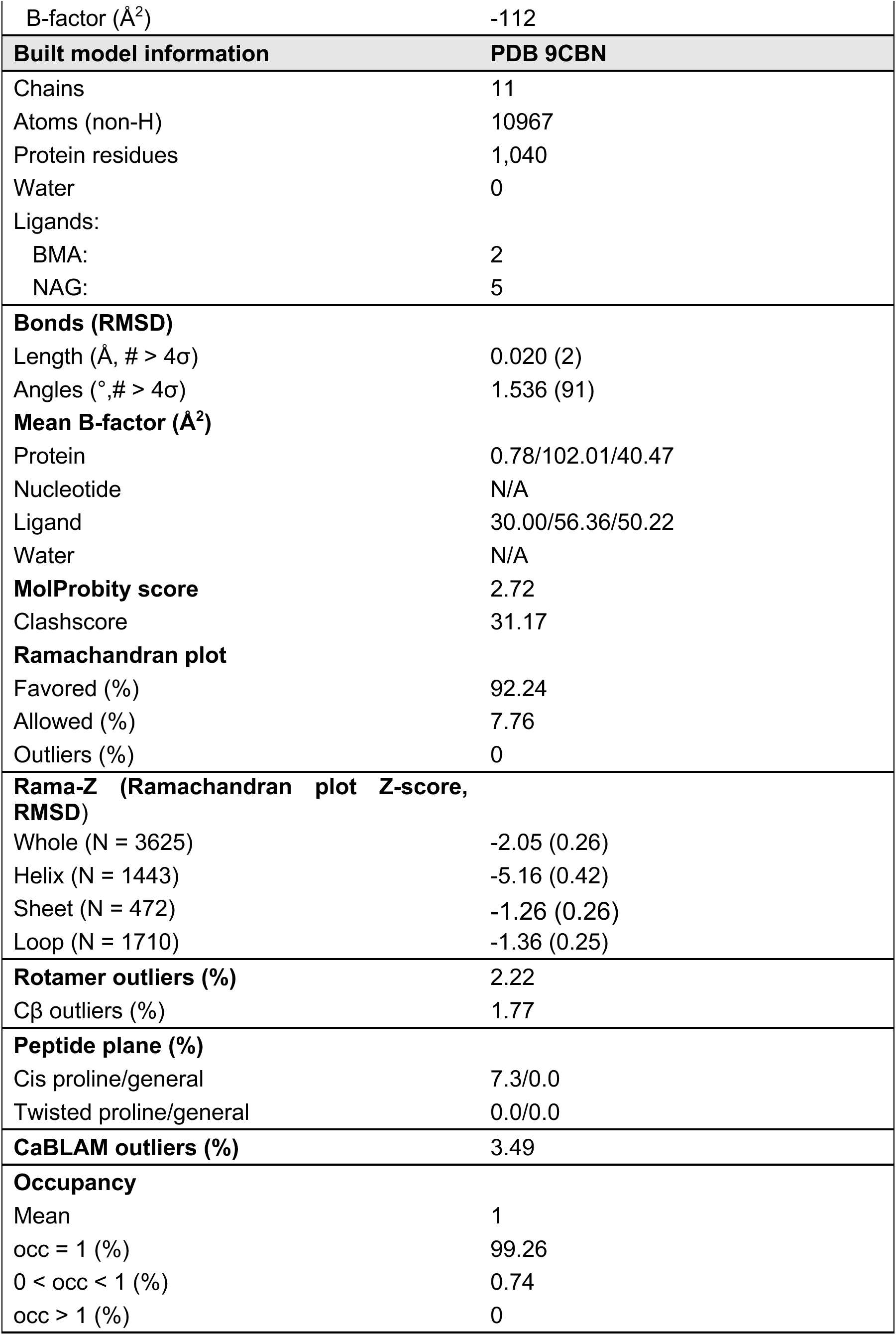
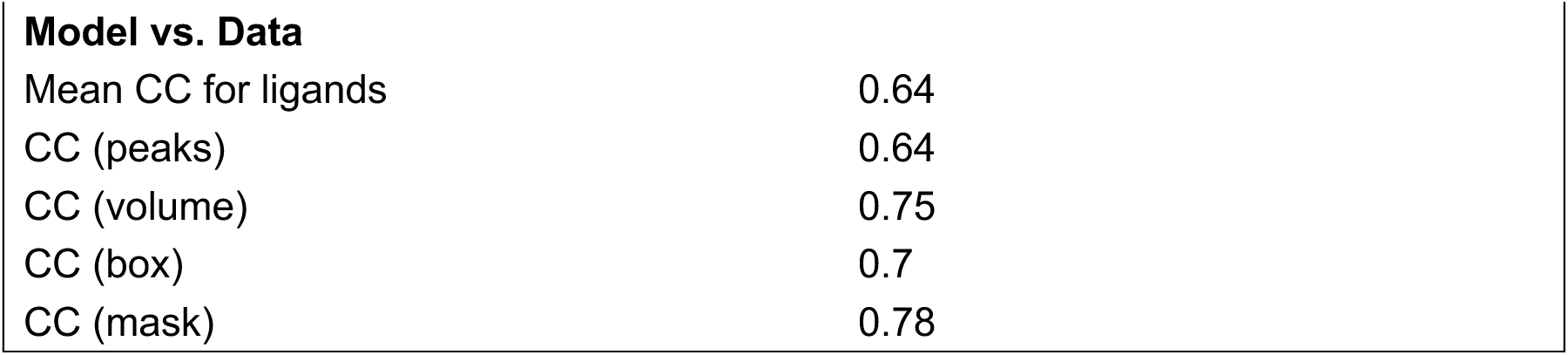
Statistics for cryoEM structure of Fab 3B4 / Fab 3H4 / HAstV1 spike complex.

| Data collection information | PNCC #160258 – Krios-3 |
| --- | --- |
| Nominal magnification | 105,000x |
| Voltage (kV) | 300 |
| Electron dose (e<sup>-</sup>/Å<sup>2</sup>) | 32.26 |
| Physical pixel size (super-res) (Å) | 0.415 |
| Movies amount | 7,235 |
| Defocus average and range (µm) | -1.5 (-2.5 to -0.5) |
| Frames | 60 |
| Single-particle reconstruction information | EMDB 45427 |
| Initial particles picked | 4,132,753 |
| Particles in the final map | 163,237 |
| Initial model used | PDB 5EWO, ab-initio |
| Symmetry imposed | C1 |
| Map overall resolution - FSC<sub>0.143</sub> (Å) | 3.33 |
| Map resolution range (Å) | 3.0-4.2 |
| Map sharpening |  |
| B-factor ($\text{\AA}^2$) | -112 |
| **Built model information** | **PDB 9CBN** |
| Chains | 11 |
| Atoms (non-H) | 10967 |
| Protein residues | 1,040 |
| Water | 0 |
| Ligands: |  |
| BMA: | 2 |
| NAG: | 5 |
| **Bonds (RMSD)** |  |
| Length ($\text{\AA}$, # > $4\sigma$) | 0.020 (2) |
| Angles ($^\circ$, # > $4\sigma$) | 1.536 (91) |
| **Mean B-factor ($\text{\AA}^2$)** |  |
| Protein | 0.78/102.01/40.47 |
| Nucleotide | N/A |
| Ligand | 30.00/56.36/50.22 |
| Water | N/A |
| **MolProbity score** | 2.72 |
| Clashscore | 31.17 |
| **Ramachandran plot** |  |
| Favored (%) | 92.24 |
| Allowed (%) | 7.76 |
| Outliers (%) | 0 |
| **Rama-Z (Ramachandran plot Z-score, RMSD)** |  |
| Whole (N = 3625) | -2.05 (0.26) |
| Helix (N = 1443) | -5.16 (0.42) |
| Sheet (N = 472) | -1.26 (0.26) |
| Loop (N = 1710) | -1.36 (0.25) |
| **Rotamer outliers (%)** | 2.22 |
| Cβ outliers (%) | 1.77 |
| **Peptide plane (%)** |  |
| Cis proline/general | 7.3/0.0 |
| Twisted proline/general | 0.0/0.0 |
| **CaBLAM outliers (%)** | 3.49 |
| **Occupancy** |  |
| Mean | 1 |
| occ = 1 (%) | 99.26 |
| 0 < occ < 1 (%) | 0.74 |
| occ > 1 (%) | 0 |
| **Model vs. Data** |  |
| Mean CC for ligands | 0.64 |
| CC (peaks) | 0.64 |
| CC (volume) | 0.75 |
| CC (box) | 0.7 |
| CC (mask) | 0.78 |

### HAstV2-neutralizing antibody binds to a distinct epitope on the upper loops of HAstV2 spike

Currently, the epitopes for only three HAstV-neutralizing monoclonal antibodies have been structurally defined, which revealed two immunogenic sites on the spike.^29,30^ Subsequently, we sought to characterize three additional neutralizing antibodies, 3B4, 3H4, and 4B6, to determine if other immunogenic sites on the HAstV spike exist.

Previous escape mutation studies identified two adjacent amino acid changes in the HAstV2 spike, D564E and N565D, which allowed HAstV2 to overcome the neutralizing activity of antibody 4B6.^26^ However, the epitope of mAb 4B6 has not been structurally defined. To visualize where neutralizing antibody 4B6 binds to the HAstV spike, we solved the crystal structure of the recombinant single-chain variable fragment (scFv) 4B6 in complex with the HAstV2 spike to 2.67 Å resolution (Fig. 3). This structure revealed that 4B6 binds to a novel 694 Å^2^ quaternary epitope at the top of the spike.

**Figure 3:**
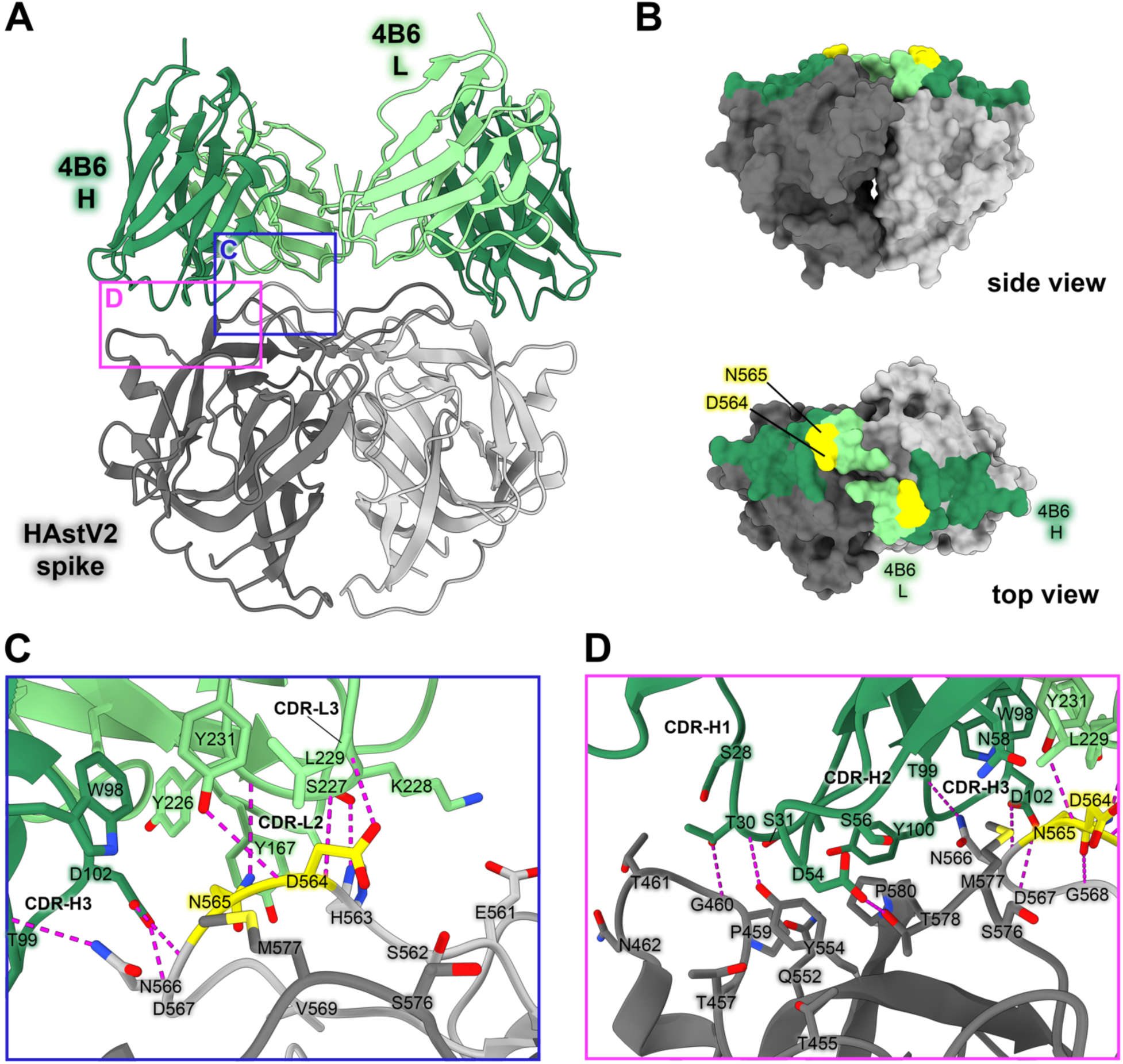
Neutralizing antibody 4B6 binds to a unique epitope on the top of the HAstV2 spike. (A) Crystal structure of scFv 4B6 bound to the HAstV2 spike homodimer solved to 2.67 Å resolution and displayed as a ribbon model. The spike is colored grey and scFv 4B6 is colored green, with the heavy chain colored dark green and the light chain colored light green. Red panels show the locations of the focused views shown in panels C and D. (B) Surface model of the HAstV2 spike with residues involved in the 4B6 epitope colored in dark green for a heavy chain interactions or light green for light chain interactions. The yellow residues indicate previously identified escape mutation locations to antibody 4B6.^12^ (C) Focused view on the light chain interaction, with 4B6 light chain colored light green. Side chains involved in hydrogen bonding are shown, with hydrogen bonds colored magenta. 4B6 light chain predominantly interacts with spike loop 3. (D) Focused view on the heavy chain interaction, with 4B6 heavy chain colored dark green. 4B6 heavy chain predominantly interacts with beta sheets 8 and 11, and the tip of loop 3 on the HAstV spike.

Each chain of 4B6 interacts predominantly with the long loop 3 from the opposing protomer, with some residues in the CDR-H3 loop of the heavy chain interacting with both protomers (Fig. 3C,D). All 3 CDR’s in 4B6 heavy chain interact with the spike, but in the light chain, only CDR-L1 and CDR-L3 interact. Antibody 4B6 forms a network of 8 hydrogen bonds with spike residues 563-567 at the very tip of loop 3, which interact with both light chain CDR L3 residues Y226-Y231 and heavy chain CDR H3 residues D102 and T99 (Fig. 3C). This hydrogen bond network consists of a mix of side-chain and backbone interactions for both the antibody and spike. This data correlates with the two residues D564 and N565 on loop 3 that were previously identified as locations for escape mutations to antibody 4B6—the mutation of these two residues would disrupt at least 2 hydrogen bond interactions, which may explain how these escape mutations disrupt 4B6 neutralization of HAstV (Fig. 3B). The HAstV spike loop 3, which 4B6 primarily targets, is highly variable across strains of HAstV, which may indicate that this location is particularly immunogenic and frequently targeted by antibodies, creating selective pressure for the virus to mutate this region.

HAstV1-neutralizing antibody 3H4 binds to a novel epitope near the base of the spike, and HAstV1-neutralizing antibody 3B4 binds the top dimer interface in a unique asymmetric way.

Previous escape mutation studies revealed a single point mutation K504E (for 3H4) or S560P (for 3B4) in the HAstV1 spike that allowed the virus to escape the neutralizing effects of antibody 3H4 or 3B4. To define the full epitopes of antibodies 3H4 and 3B4, we solved the structure of both Fabs 3H4 and 3B4 in complex with the HAstV1 spike to 3.33 Å resolution using single-particle cryoEM (Fig. 4A,B, Supp. Fig. 2). This structure reveals two novel epitopes, with a single Fab 3B4 bound to the top of the spike dimer interface, and two Fab 3H4 bound to the bottom sides of the spike dimer (Fig. 4A,B). Antibody 3B4 spans a 1039 Å^2^ quaternary epitope across the top dimer interface, with more of the epitope located on one protomer than the other (Fig. 4A,E). Based on the structure, as well as the retention volume of the complex in solution on a size-exclusion chromatography column, only one Fab 3B4 can bind the spike homodimer at a time, which represents the first antibody of its kind to be discovered for HAstV, as all other previously characterized antibodies can bind symmetrically with one antibody binding site per protomer. Antibody 3B4 targets the majority of loop 3 on one monomer closer to the base of the loop, and the side of loop 3 on the other monomer. Although 3B4 targets a similar structural region on HAstV1 spike as that of 4B6 on HAstV2 spike, it interacts with unique residues focused more on the center of the dimer interface, while 4B6 is targeted more outward towards the tips of loop 3. All six of 3B4 CDR loops interact with the spike, forming a hydrogen bond network primarily between spike residues G573- T577, and 3B4 residues S30-N32 on CDR-L1 (Fig. 4C). Q53 and S50 from CDR-L2 also contribute several hydrogen bonds, with Q53 making two hydrogen bonds with T613 and N614. On the heavy chain, the majority of the hydrogen bonds are contributed by CDR-H1 residues T28 and T30, which target T562 and S560 on the spike. Although a single point mutation of S560P in the spike sequence confers resistance to antibody 3B4 neutralization, this mutation actually changes two distinct sites of interaction with the heavy chain of 3B4, given the close locations of each S560 to each other on the dimer interface (Fig. 4C,E). This could suggest that single point mutations offer higher resistance to dimer interface antibodies in comparison to antibodies that bind both protomers. Despite both 4B6 and 3B4 targeting loop 3, 3B4 targets residues that are more conserved.

**Figure 4:**
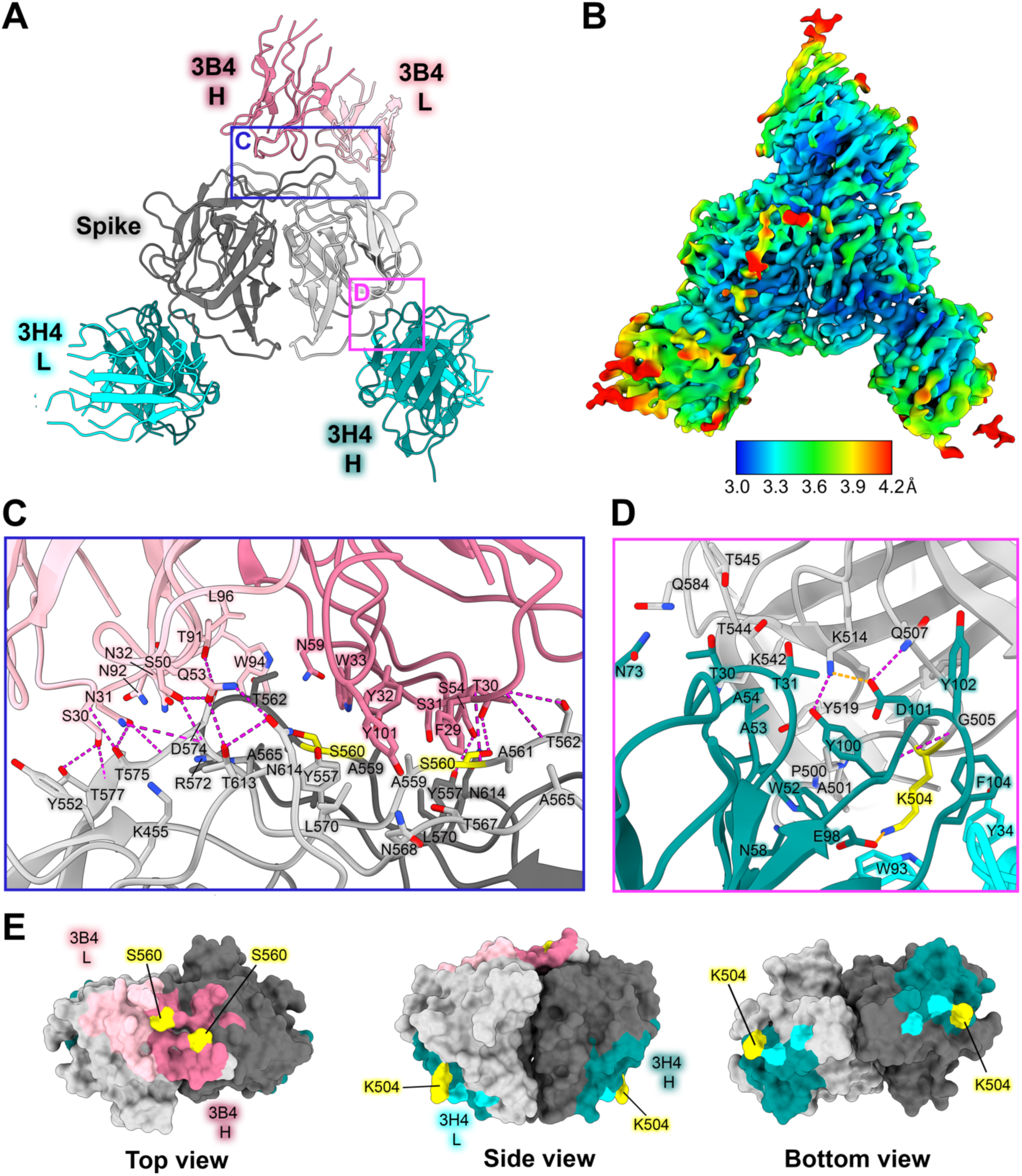
Neutralizing antibody 3H4 binds to a unique epitope at the base of the spike, and neutralizing antibody 3B4 has a unique top epitope in which a single antibody binds the spike dimer interface. (A) Single-particle cryoEM reconstructed map solved to FSC_0.143_ 3.33 Å of neutralizing Fab 3H4 and Fab 3B4 bound simultaneously to the HAstV1 spike, displayed as a ribbon model with 3H4 colored cyan and 3B4 colored pink. The heavy and light chains are colored in dark and light shades, respectively. Red panels show the locations of the focused views shown in panel C and D. (B) Local resolution estimation of the cryoEM structure of HAstV1 spike bound to 3H4 Fab and 3B4 Fab, with contour level at 0.043 in ChimeraX. (C) Focused view of the 3B4 epitope, with the light chain colored light pink, and the heavy chain colored dark pink, with hydrogen bond interactions colored magenta. Serine 560, which was previously identified as a residue that overcomes the neutralization activity of 3B4 when mutated to proline, is highlighted in yellow. (D) Focused view of the 3H4 epitope, with the light chain colored light cyan, and the heavy chain colored dark teal. Hydrogen bond interactions are colored magenta and salt bridges are colored in orange. Lysine 504, which was previously identified as a residue that overcomes the neutralization activity of 3H4 when mutated to glutamic acid, is highlighted in yellow. (E) Surface view of the HAstV1 spike with antibody interacting residues colored according to antibody chain. Residues interacting with both chains are colored according to the predominant interaction. Residues that confer resistance to the respective antibody when mutated are colored in yellow.

Fab 3H4 binds to a novel 676 Å^2^ epitope near the base of the spike which is distinct from any other known HAstV-neutralizing antibody epitopes, as all previously solved spike-antibody structures target the top or upper sides of the spike dimer (Fig. 4A,B). The 3H4 epitope interaction is facilitated almost entirely by the heavy chain alone, with only W93 from CDR L1 and Y34 CDR L3 from the light chain making any contact with the spike (Fig. 4D,E). Antibody 3H4 mostly targets the upper portion of the spike loop 2 with all 3 heavy chain CDR loops. Two salt bridges formed between K514 on spike and D101 on CDR-H3, and K504 with E98 on CDR-H3 (Fig. 4D). These lysine residues also form hydrogen bonds and cation-pi interactions with Fab 3H4. Notably, the salt bridge interaction by K504 appears critical to the ability of 3H4 to bind to spike as the mutation of K504 to a negatively-charged glutamic acid disrupts 3H4 neutralization of HAstV1.^26^ Antibody 3H4 also targets a region of much higher conservation than that of the other antibodies, with over 70% of the interacting spike residues being semi-conserved or higher amongst the 8 HAstV serotypes. Despite the majority of residues being conserved, K504, which is critical to 3H4 neutralization, is highly variable among serotypes, which likely accounts for the 3H4 serotype specificity.^26^

### 3B4, 3H4, and 4B6 scFv neutralize HAstV

Antibody 3H4 reveals a particularly interesting epitope location, as the full-length antibody would likely clash with the icosahedral core of the HAstV capsid, suggesting that this antibody may contort the spike dimer in some way. Since 3H4 binds so distantly from other structurally determined neutralizing antibody epitopes and yet is still shown to block FcRn receptor binding, we hypothesized that 3H4 may neutralize HAstV by steric hinderance with its constant regions and contortion of the spike, rather than the direct blocking of an important functional site on the spike. We tested whether scFv 3H4, scFv 3B4, and scFv 4B6 which lack antibody constant domains, could still neutralize HAstV1 (scFv 3H4 and scFv 3B4) or HAstV2 (scFv 4B6). HAstV1 was preincubated with serial dilutions of scFv 3H4, scFv 3B4, or mAb 3B4 as a control, or HAstV2 with scFv 4B6, and was incubated on a Caco-2 cell monolayer, and viral infection was measured by an immunoperoxidase focus-forming assay. We found that both the scFv 3H4, 3B4, and 4B6 are still able to neutralize HAstV and do so to a similar degree, but are not as effective at neutralization as full-length mAb (Fig. 5B), indicating that steric hinderance and/or avidity have a role in the ability of these antibodies to neutralize virus. 4B6 appears to be the most affective at neutralization despite its lower affinity, however, this could also be a function of it neutralizing a different serotype than 3H4 and 3B4.

**Figure 5:**
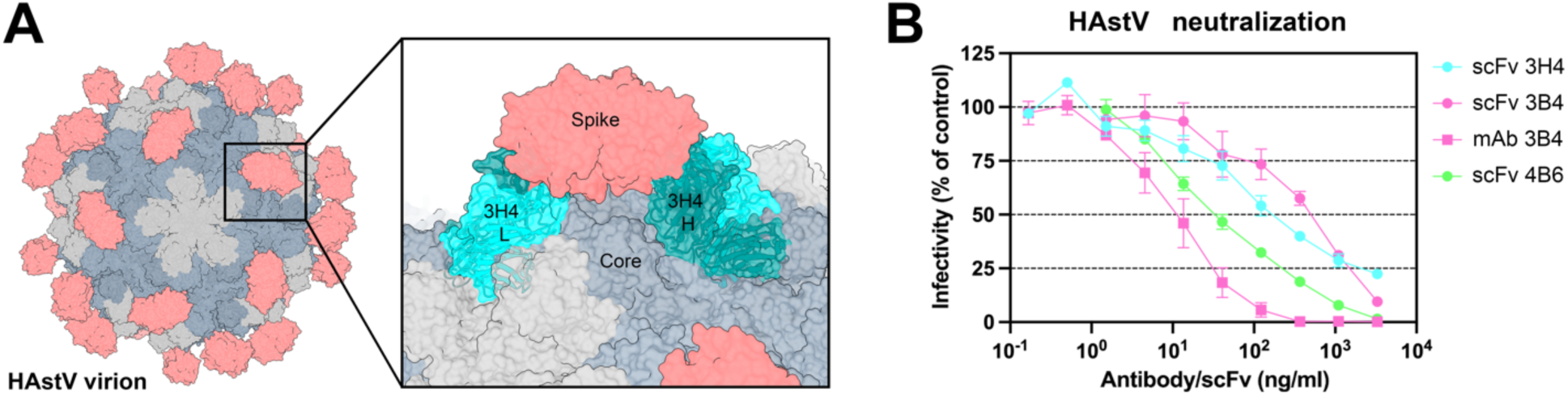
Steric hinderance of antibody 3H4 constant domains may play a role it its ability to neutralize HAstV1. (A) Graphic depicting the full virion capsid, with the core domains colored in grey and the spike domains colored in salmon. The panel shows a focused view of how Fab 3H4 would clash with the HAstV capsid core, using the cryoEM reconstruction of 3H4 variable domain aligned with an AlphaFold 3 model of the constant domain. (B) Neutralization activity of scFv 3H4, scFv 3B4 and mAb 3B4 against HAstV1, or scFv 4B6 against HAstV2. HAstV was preincubated with the corresponding scFv or mAb at the indicated concentrations. The infectivity of the virus was determined as described in Materials and Methods. The infectivity assay was performed in biological triplicates and carried out in duplicate. The data are expressed as % infectivity of control and represent the mean ± SEM.

### AlphaFold3 prediction accuracy

As predictive protein structural software advances, we sought to assess how accurately the recent release of AlphaFold 3 (AF3) could predict antibody-antigen interactions.^39^ We compared the crystal structure of scFv 4B6 and HAstV2 spike with that of its AlphaFold prediction, and found that not only was the antibody placed correctly, but even the side chain interactions were highly accurate (Supp. Fig. 3A). However, on a macroscopic scale, the dimer interfaces appear to be slightly misaligned, causing the other protomer alignment and subsequent interacting residues to be slightly misaligned (Supp. Fig. 3D,E), though the local side chain orientations still appear to be highly accurate. AF3 was confident in its prediction, with pTM of 0.88 and ipTM of 0.86. The overall accuracy of the AF3 model is quite high, with a TM-score of 0.97 (TM value of 1=identical match) when the AF3 model is aligned to the crystal structure. This is a substantial improvement from the AlphaFold 2 (AF2) prediction, which did not place scFv 4B6 in the correct general placement, let alone correct side chain orientations (Supp. Fig 3A). We additionally compared the AF3 model of Fab 3H4 and Fab 3B4 with our solved cryoEM structure. The AF3 model of Fab 3H4 bound to HAstV1 spike was highly accurate (Supp. Fig.3B), with TM-score of 0.99 when aligned to the cryoEM structure with Fab 3B4 removed. Despite the higher TM-score, AF3 reported slightly lower confidence scores, with ipTM=0.78, and pTM=0.81. The AF3 predicted model for 3H4 additionally showed dramatic improvement from the AlphaFold 2 (AF2) prediction, which did not place Fab 3H4 in the correct general placement. In the case of Fab 3B4, AF3 could not successfully find the correct general placement and consistently placed 3B4 Fab on the side of the spike dimer (Supp.Fig.3C), even when we tried alternative searches for one or two Fabs or scFvs. Because the overall interface alignment appears to be slightly off in these AF3 models, this may explain why AF3 could not predict the epitope of Fab 3B4 correctly, which targets the dimer interface. AF3 was less confident in its predicted model of one Fab 3B4 bound to HAstV1 spike, with ipTM=0.52 and pTM=0.61, but were still above the 0.5 threshold suggesting that the structure could be correct despite being an incorrect placement. However, the decrease in these scores for 3B4 compared to 3H4 and 4B6 does suggest some ability to determine whether the predicted structure is correct. From these assessments, AlphaFold 3 appears to have a dramatic increase in accuracy compared to previous versions which consistently failed to predict antibody interactions at all, even though some challenging antibodies which target interfaces may still be more difficult.

## Discussion

Here, we map three new epitopes on the HAstV spike that induce neutralizing antibodies, finding that 4B6 and 3B4 target the top of the spike in ways that are unique from previously characterized antibodies. Additionally, we find that 3H4 targets the base of the spike, representing an entirely unique epitope which is distant from previously characterized antibodies and targets mostly conserved residues. With these additional structures, we find that the majority of neutralizing antibodies target the upper side or top variable loop regions of the spike (Fig. 6). These regions reside around conserved areas of the HAstV spike, termed the P-site and S-site, which were proposed as potential host protein interacting.^40^ However, few of the antibody residues directly target these conserved sites. Although the direct receptor or host protein interaction locations on the spike are currently not known, it is possible that these neutralizing antibodies sterically hinder receptor binding to more conserved residues, rather than overlapping with receptor binding site(s) directly, given that the majority of neutralizing antibodies target highly variable loop residues on the top regions of the spike, indicating less functional importance for these residues. These loops may serve more as an immunogenic target for antibodies that can be more easily mutated without changing important functions of the spike. 3H4 represents an entirely new antigenic site near the base of the spike, which has less accessibility compared to the top exposed portion of the spike where the majority of antibodies target. This low epitope may have been favored more by recombinant spike immunization than what would have been induced by the whole virus where the capsid core domain limits access. This suggests the possibility of using recombinant spike vaccinations for enhancing the induction of less accessible antigenic sites that may be more conserved on the spike, similar to how some recombinant influenza vaccine antigen candidates better elicit immune- subdominant hemagglutinin stem targeting antibodies. Given that 3H4 targets mostly conserved residues and has high affinity, it may have some potential as a monoclonal antibody therapy for HAstV2. However, 3H4 is vulnerable to mutations at residue K504, which additionally is not conserved between HAstV serotypes.

**Figure 6:**
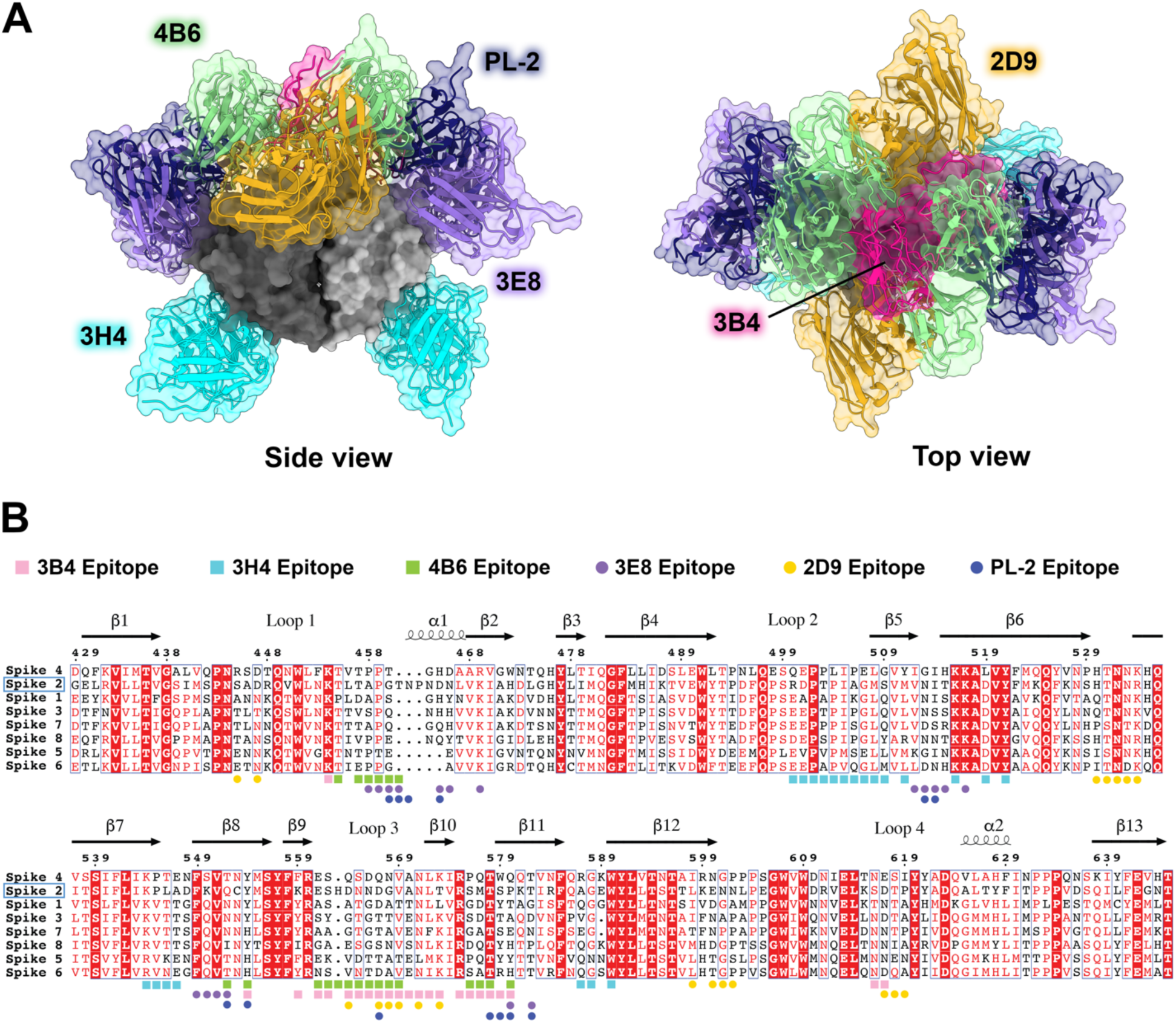
Comparison of all known HAstV-neutralizing antibody epitopes, showing that most target the upper variable region of HAstV spike. (A) Alignment of all existing HAstV neutralizing antibody structures 4B6, 3B4, 3H4, 3E8, 2D9, and PL- 2, mapped onto HAstV1 spike. (B) Spike protein sequences of the eight classical HAstV serotypes aligned using EMBL-MUSCLE, with residues colored according to conservation. The following sequences were used for the alignment: HAstV1, GenBank #AAC34717.1; HAstV2, GenBank # KY964327.1; HAstV3, UniProt #Q9WFZ0.1; HAstV4, UniProt #Q3ZN05.1; HAstV5, UniProt #Q4TWH7.1; HAstV6, UniProt #Q67815.1; HAstV7, UniProt #Q96818.2; HAstV8, UniProt #Q9IFX1.2. Residue numbering shown above corresponds with HAstV2. Residues highlighted in red are strictly conserved, residues with red text are semi-conserved, and residues in black text have little to no conservation. Spike residues interacting with the antibodies characterized in this paper, 3H4, 3B4, and 4B6, are indicted with colored squares, and epitope residues for antibodies that were previously characterized, 2D9, 3E8, and PL-2, are indicated as colored circles.

We find that all three antibodies, 3H4, 4B6, and 3B4, block binding of FcRn to the spike, although 3B4 only appears to partially block. From the structure, is can be seen that 3B4 leans more to one side of the spike dimer than the other. It is possible that this asymmetric nature of the 3B4 binding antibody could explain how only partial blocking of FcRn binding occurs if FcRn were to bind both sides of the spike homodimer and 3B4 was capable of only blocking one side. It is interesting that all three antibodies block FcRn binding given their different locations on the spike, which leads to our hypothesis that the blocking ability of these antibodies may be more related to steric hinderance and less related to where the antibodies bind directly. This does seem to be the case given that the scFvs of 3H4 and 3B4 neutralize less effectively than full length mAb, however they are still able to neutralize virus at higher concentrations, indicating that there may still be some overlap with receptor-binding site(s), or that some steric hinderance still occurs with the variable region.

Overall, these studies further our structural and mechanistic understanding of neutralizing antibody epitopes on the HAstV capsid surface, supporting the rational design of vaccines targeting HAstV spikes to prevent childhood viral diarrhea by HAstV.

## Acknowledgements

This research was funded by NIH grant R01 AI144090 to R.M.D. and C.F.A. This work was partially supported by M0037-Fordecyt grant 302965 from the National Council for Science and Technology-Mexico (CONAHCyT) to S. López. S. Lanning was supported by the NIH training grant T32 GM133391. Funding for the purchase of the Octet RED384 instrument was supported by the NIH S10 shared instrumentation grant 1S10OD027012-01. This research used resources of the Advanced Photon Source, a U.S. Department of Energy (DOE) Office of Science user facility operated for the DOE Office of Science by Argonne National Laboratory under Contract No. DE-AC02- 06CH11357. A portion of this research was supported by NIH grant U24GM129547 and performed at the PNCC at OHSU and accessed through EMSL (grid.436923.9), a DOE Office of Science User Facility sponsored by the Office of Biological and Environmental Research.

